# On the performance of multi-compartment relaxometry for myelin water imaging – Intra-subject and inter-protocol reproducibility

**DOI:** 10.1101/2022.09.07.506917

**Authors:** Kwok-Shing Chan, Maxime Chamberland, José P. Marques

**Affiliations:** Donders Institute for Brain, Cognition and Behaviour, Radboud University, Nijmegen, The Netherlands

**Keywords:** Myelin water imaging, diffusion weighted imaging, microstructure, gradient echo imaging

## Abstract

We evaluate the test-retest repeatability and study the tissue properties of multicompartment relaxometry-based myelin water imaging (MCR-MWI) derived from different gradient echo (GRE) acquisition settings. Additionally, the variable flip angle acquisition scheme is optimised based on numerical simulations to reduce the acquisition time of MCR-MWI in a clinically practical range without using advanced image acquisition methods. For the test-retest analysis, in vivo imaging was performed to collect data from three healthy volunteers in two identical sessions. Three GRE sequence settings with different echo times and repetition times imitating various scanner setups were evaluated. The in vivo data was also used to validate the optimal variable flip angle combination derived from simulations. Bundle-specific profiles of MCR-MWI derived microstructural parameters were investigated, as well as the cross-correlations of those parameters. Good cross-session repeatability is observed for MCR-MWI. While good correlations can also be found in myelin water fraction (MWF) across protocols, systematic differences, particularly for protocols with different repetition times, are observed. Numerical simulations indicate that MCR-MWI can be performed with a minimum of three flip angles covering a wide range of T_1_ weighting without adding significant measurement bias and the result is supported by the in vivo experiment allowing whole brain 1.5mm isotropic MWF maps to be acquired in 9 minutes. Bundles-specific MWF analysis reveals that certain white matter bundles are similar in all three participants. We also found that microstructure relaxation parameters have low correlations with MWF. MCR-MWI is a reproducible measure of myelin. However, attention should be paid to considering the protocol related MWF differences for comparison studies, especially when different repetition times are used as this can introduce biases up to 0.5% of MWF in our tested protocols. The optimised flip angle acquisition scheme can reduce the total scan time to 40% of the original implementation without significant quality degradation.

**Highlights:**

- Multi-compartment relaxometry based myelin water imaging (MCR-MWI) can be performed with data comprising as few as 3 flip angles without introducing substantial bias or instability in the fitting procedure;
- MCR-MWI is a reproducible measure of myelin water fraction (MWF) and incorporating DWI can further improve the measurement reproducibility;
- MCR-MWI allows the acquisition of whole brain 1.5mm isotropic MWF maps in 9 minutes, even without the use of advanced model-based reconstructions;
- Small MWF bias can present in cross-protocol comparison if the MT effect is not constant across GRE protocols (e.g., different TRs or flip angle combinations);
- Compartmental relaxation parameters derived from MCR-MWI possess complimentary information beyond myelin water concentration.

## 1. Introduction

Myelin water imaging (MWI) using multi-echo gradient echo (mGRE) acquisition to probe myelin in white matter (WM) has been recently introduced and received increasing attention because of its simple implementation and high scan time efficiency (Nam et al., 2015b). However, data fitting with a 3-pool exponential model to the GRE signal is an ill-conditioned problem (Lee et al., 2016). To overcome the challenges facing the conventional GRE-MWI approach, multi-compartment relaxometry for MWI (MCR-MWI) was introduced (Chan and Marques, 2020). Briefly, myelin water fraction (MWF) measurement can benefit from a variable flip angle (VFA) acquisition strategy, simultaneously exploiting the apparent transverse and longitudinal relaxation differences between free (intra- and extra-axonal) water and myelin water as well as their different frequency shifts. As a result, MCR-MWI not only corrects the T_1_-saturation dependence on the observed MWF but also improves the fitting condition when solving the ill-posed MWI problem. MCR-MWI can work with multi-shell diffusion-weighted imaging (DWI) data and the hollow cylinder biophysical fibre model (MCR-DIMWI) to further improve the SNR of the MWF estimation.

While MCR-MWI demonstrated high sensitivity and specificity to myelin water, some criteria must be met before the method can be applied to a broader range of scientific and clinical applications. One of them is that the method must be performed in a clinically feasible scan time (considering similar advanced methodologies we assume it to be in the range of 10 to 15 min). In the initial implementation, we utilised 7 different flip angles ranging from 5° to 70° in the VFA acquisition to ensure a diversity of T_1_-weighting was added to the signal, yet unavoidably increased the scan duration, albeit comparable to a previous study using the conventional GRE-MWI method (Nam et al., 2015b). Given that the high-quality MWF results have been established with MCR-MWI (Chan and Marques, 2020), reducing the number of flip angles in the acquisition with further optimisation on the usage of flip angle combination can potentially reduce the scan time without compromising its robustness. Another crucial factor for longitudinal studies or multi-centre trials is that the method must be reproducible, both within-subjects and across acquisition protocols.

In this study, we set out to investigate if the acquisition time for MCR-MWI could be shortened by reducing the number of flip angles without compromising its robustness. In the second part, we conduct a test-retest experiment to evaluate the cross-protocol repeatability of MCR-MWI and MCR-DIMWI, using 3 sets of acquisition protocols with different echo spacing and repetition times. Additionally, we also study the reproducibility of some of the MWI-derived tissue properties across WM fibre bundles.

## 2. Material and Methods

### 2.1. Numerical Simulations for scan time optimisation

Monte Carlo simulations were conducted to investigate the impact on the MWF estimation robustness when different numbers of flip angles are used in data acquisition, using the MCR-DIMWI signal model to generate a 3-pool WM signal (tissue parameters and acquisition protocol settings shown in Supplementary Material). For each flip angle, the same magnitude of Gaussian noise was introduced to the simulated signal to achieve moderate SNR (SNR_TE0_=57 when *α*=20°). The data were then fitted to the MCR-MWI signal model using either the full set of 7 flip angles or a subset of the 7 flip angles that comprised data with only 2 to 5 flip angles. The process was repeated 4000 times such that statistics can be derived to compare the performance of different flip angle choices.

MWF measurement accuracy and precision were assessed by examining the bias and standard deviation (SD) of the fitted MWF to the ground truth:

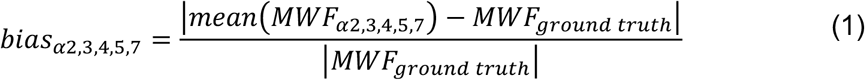

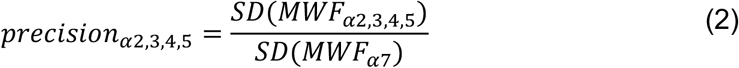

where MWF_ground truth_ is the ground truth MWF in the simulated signal, mean(MWF_*α*2,3,4,5_) is the mean of the estimated MWF for each flip angle subset (of 2,3,4 or 5 flip angles respectively) across the 4000 trials and MWF_*α*7_ is the estimated MWF using the full set of the 7 flip angles. If the computed bias and precision were lower than the 7-flip-angle protocol (considered as state-of-the-art) they were thresholded at bias_*α7*_ and 1. An ad hoc performance metric was derived to determine the optimum subset of flip angle combination that can yield a similar performance as using the full dataset:

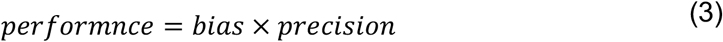

A value of the performance close to 1 indicates that the subset performs as good as the full dataset, as its value increases the performance is deteriorating.

### 2.2. In vivo imaging

Data acquisition was performed at 3T (Prisma, Erlangen, Siemens) on 3 healthy volunteers (age 30, 43 and 61 years, all males). Test-retest repeatability of MCR-(DI)MWI was evaluated by acquiring imaging data in 3 experiment scenarios which were repeated in 2 separate imaging sessions, corresponding to:

1. **Short scan time condition (Prot1):** TR/TE_1_/ΔTE/#TE = 38/2.2/3.07 ms/12, TA = 2.8 min per flip angle;
2. **Data-rich condition (Prot2):** TR/TE_1_/ΔTE/#TE = 50/2.2/3.07 ms/15, TA = 3.5 min per flip angle; and
3. **Less-performing gradient setup (Prot3):** TR/TE_1_/ΔTE/#TE = 55/2.68/3.95 ms/13, TA = 4 min per flip angle,

with sagittal slice orientation, FOV = 168×222×240 mm^3^, resolution = 1.5 mm isotropic, *α* = [5, 10, 20, 50, 70]°, a 5-fold acquisition acceleration was used based on CAIPI under-sampling (CAIPI z-shift of 2). B_1_ map was acquired using a turbo-flash protocol to correct B_1_ field inhomogeneities on the VFA-mGRE data (Chung et al., 2010).

DWI data was acquired in one of the sessions for MCR-DIMWI and bundle-specific analysis with parameters: 2D-MB EPI-DWI, MB = 3, res = 1.6 mm isotropic, TR/TE = 3350/71.20 ms, two shells (b=0/1250/2500 s/mm^2^, 17/120/120 diffusion-encoding directions), TA = 15 min. A summary of the experiment design is illustrated in Figure 1.

**Figure 1:**
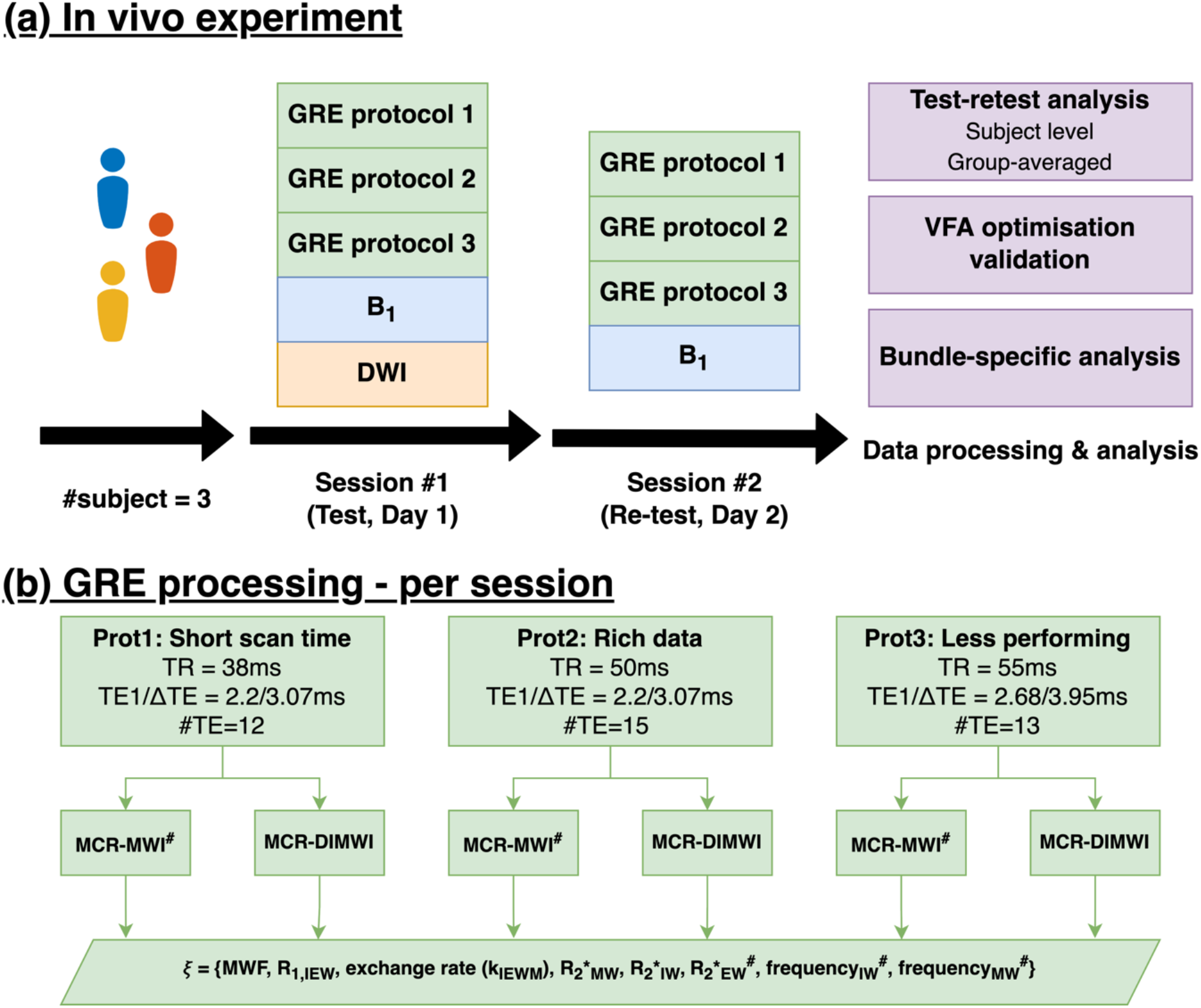
A summary of the experimental design. (a) Data acquisition was conducted in two separate sessions. Each of them consists of 3 VFA-mGRE protocols with different sequence settings. (b) The major sequence parameter differences between the 3 GRE protocols. Data from each protocol was fitted independently with the MCR-MWI and MCR-DIMWI models to derive MWF as well as other microstructural parameters. ^#^Parameters available only for MCR-MWI

### 2.3. Data processing

Image registration was first performed within each GRE protocol to mitigate minor head movements across the GRE acquisitions using rigid body transform with linear interpolation (Avants et al., 2011). B_1_ maps were then registered to the GRE data. Voxel-wise fitting was then performed with the MCR-(DI)MWI models to obtain MWF and other tissue properties, corresponding to the following optimisation problem:

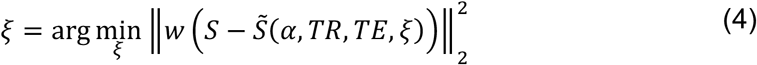

where *S* is the measured signal and 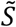 is the simulated signal using the MCR-(DI)MWI model given the compartmental tissue parameter set *ξ* and the acquisition parameters, including flip angle (*α*), repetition time (TR) and echo time (TE), and *w* is a weighting term. Based on our observation of the data quality and the preliminary analysis on various weighting methods (see Appendix A), a quadratic signal weighting strategy was used in this study for the model fitting. Specifically, for data acquired at TE and *α*,

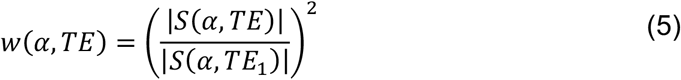

Data of all 5 flip angles for Scenario 1 (short scan time protocol, FA5-Prot1), Scenario 2 (rich data protocol, FA5-Prot2) and Scenario 3 (less performing gradient protocol, FA5-Prot3) were used in both MCR-MWI and MCR-DIMWI to derive the tissue parameters. Validation of the flip angle optimisation findings from numerical simulations was performed using only 3 flip angles from the full dataset. Four flip angle combinations were selected based on the performance score from Eq. (3), including the top 3 performers and the one with the poorest score. The flip angle combinations chosen were: [10,20,70]° (referred to as FA3[10,20,70]), [5,10,70]° (FA3[5,10,70]), [10,20,50]° (FA3[10,20,50]) and [5,10,20]° (FA3[Poor]). All MWI results were subsequently registered to the DWI space using rigid body transform with linear interpolation for statistical analysis.

DWI data was pre-processed using an in-house pipeline consisting of the following procedures: (1) data denoising with Patch2self (Fadnavis et al., 2020; Garyfallidis et al., 2014), (2) Gibbs ringing removal (Kellner et al., 2016; Tournier et al., 2019), (3) susceptibility distortion correction (Andersson et al., 2003; Smith et al., 2004) and (4) eddy current distortion correction (Andersson and Sotiropoulos, 2016). Spherical mean technique (SMT) (Kaden et al., 2016) and a ball-and-stick model (Behrens et al., 2007) were used to compute WM properties such as intra-axonal volume fraction and fibre orientations for MCR-DIMWI.

Bundle-specific analysis was used in the statistical analysis, with 50 subject-specific WM bundles obtained from TractSeg (Wasserthal et al., 2018), using multi-shell, multi-tissue constrained spherical deconvolution (Jeurissen et al., 2014). Tractometry (Chamberland et al., 2021; Wasserthal et al., 2020) was performed to compute the bundle-specific profiles on MWF and other metrics at 100 locations along each tract. To avoid the increasing partial volume effect toward the ends of the bundles, only 98 sections in the middle were used, resulting in 4900 WM region of interests (ROIs) per subject.

### 2.4. Data analysis

#### 2.4.1. MWF test-retest repeatability

Test-retest reliability was assessed via Pearson’s correlation coefficient (R) and coefficient of variation (CV, standard deviation of the differences divided by the mean) for each GRE protocol (same protocol, cross-session). The same metrics were also derived when comparing the results between various protocols (cross-protocol with data from the two sessions jointly analysed). The analysis was performed at two levels: (1) at the subject level (4900 ROIs for each subject) and (2) using the WM ROIs from all three subjects together (14700 ROIs). To be able to evaluate MWF biases due to the protocol differences, the mean MWF across the three GRE protocols and 2 imaging sessions using the data comprising all 5 flip angles was computed and defined as the MWF ground truth (GT) of each ROI. The MWF estimation bias is subsequently derived as the difference between the GT and the MWF of each protocol obtained from one of the imaging sessions.

#### 2.4.2. Validation of flip angle combination

Bland-Altman analysis was used in analysing the agreement between the MWF estimation of using the full dataset of 5 flip angles and using only 3 flip angle data of Protocol 1 (i.e., FA5-Prot1 vs FA3[10,20,70], FA5-Prot1 vs FA3[5,10,70], FA5-Prot1 vs FA3[10,20,50], and FA5-Prot1 vs FA3[Poor]) for both imaging sessions, using all 13500 ROIs of the three subjects). Since most of the fitted parameter does not exhibit a Gaussian distribution (Jarque-Bera tests rejected the null hypothesis that the inputs come from a normal distribution), the median of the MWF differences between the usage of the two datasets is reported instead of the mean. Additionally, the reproducibility coefficient (RPC) was computed from the interquartile range (IQR), i.e., 1.45×IQR.

#### 2.4.3. Bundle-specific analysis

Bundle-specific MWF profiles were visually compared on 8 major fibre bundles covering 3 main axes of the brain (left-right, superior-inferior and anterior-posterior) between MCR-MWI and MCR-DIMWI, and between subjects using the FA5-Prot2 results. These bundles are the splenium/genu of the corpus callosum (CC), the left/right optic radiation (OR), the left/right inferior longitudinal fascicles (ILF) and the left/right corticospinal tracts (CST). For all these bundles, the original TractSeg segmentation was used except for the splenium of the CC, which was filtered manually due to significant irregularities of tract terminations observed in one subject (see Supplementary Section 6).

Apart from MWF, MCR-MWI also provides free water R_1_ (R_1,IEW_), exchange rate (k_IEWM_) and intra-axonal water R_2_* maps (amongst others). Additionally, g-ration maps (ratio between inner and outer radii of the axon) can be derived using the fitted water signal amplitudes and the formalism described in (Stikov et al., 2015) and (Jung et al., 2018) as:

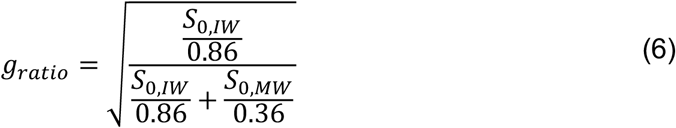

We explore the subject-averaged parameter profiles along four major bundles to investigate the potential usage of these parameters and their information independence. Protocol 2 results are also used to report the cross-correlation among these four tissue parameters.

## 3. Results

### 3.1. Optimisation and validation of flip angle combination

Optimisation of the flip angle usage for MCR-MWI based on numerical simulation is shown in Table 1. Similar accuracy compared to using data of 7 flip angles can be achieved with only 3 flip angles, whereas significant bias is observed when the number of flip angles was reduced to 2. The optimum precision progressively decreased with the amount of data available (about 1.4-fold SD of using 3 flip angle data compared to using 7 flip angles). From a data averaging perspective 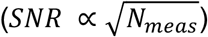, the standard deviation would be expected to decrease by 1.53-fold when using 3 rather than 7 flip angle data, suggesting that the optimised VFA acquisition schemes have increased efficiency.

**Table 1:** Top 5 subsets of the 7 flip angles in term of performance tested in the numerical simulations.

a) Using 5 flip angles
| # | Flip angle (°) | Performance | Bias | Precision |
| --- | --- | --- | --- | --- |
| 1 | [5,10,20,50,70] | 0.042 | 0.037<sup>†</sup> | 1.112 |
| 2 | [5,10,30,50,70] | 0.042 | 0.037<sup>†</sup> | 1.121 |
| 3 | [10,20,40,50,70] | 0.044 | 0.041 | 1.087 |
| 4 | [5,10,20,40,70] | 0.044 | 0.038 | 1.177 |
| 5 | [5,10,40,50,70] | 0.044 | 0.039 | 1.132 |

b) Using 4 flip angles
| # | Flip angle (°) | Performance | Bias | Precision |
| --- | --- | --- | --- | --- |
| 1 | [10,20,50,70] | 0.044 | 0.037<sup>†</sup> | 1.165 |
| 2 | [5,10,50,70] | 0.049 | 0.040 | 1.242 |
| 3 | [5,10,50,70] | 0.049 | 0.037<sup>†</sup> | 1.319 |
| 4 | [5,10,40,70] | 0.050 | 0.039 | 1.264 |
| 5 | [5,10,20,70] | 0.051 | 0.037<sup>†</sup> | 1.355 |

**c) Using 3 flip angles**
| # | Flip angle (°) | Performance | Bias | Precision |
| --- | --- | --- | --- | --- |
| 1 | [10,20,70] | 0.052 | 0.037<sup>†</sup> | 1.382 |
| 2 | [5,10,70] | 0.056 | 0.037<sup>†</sup> | 1.500 |
| 3 | [10,20,50] | 0.056 | 0.041 | 1.382 |
| 4 | [5,20,70] | 0.059 | 0.041 | 1.431 |
| 5 | [20,30,70] | 0.059 | 0.037<sup>†</sup> | 1.582 |

**d) Using 2 flip angles**
| # | Flip angle (°) | Performance | Bias | Precision |
| --- | --- | --- | --- | --- |
| 1 | [10,70] | 0.091 | 0.065 | 1.398 |
| 2 | [5,70] | 0.117 | 0.076 | 1.535 |
| 3 | [10,50] | 0.171 | 0.132 | 1.294 |
| 4 | [5,50] | 0.232 | 0.149 | 1.552 |
| 5 | [20,50] | 0.254 | 0.227 | 1.117 |

<sup>†</sup>Thresholded to the bias derived with data of 7 flip angles

Figure 2 shows the Bland-Altman analysis to validate the flip angle optimisation on in vivo MWF estimation. The best performing three flip angle combinations (FA3[10,20,70], FA3[5,10,70] and FA3[10,20,50]) only results in small bias (ranging from 0.01% to 0.63%) and low RPC (ranging from 0.83% to 1.76%) compared to using data of 5 flip angles. Additionally, the RPC of these three agrees with the simulation results where FA3[10,20,70] and FA3[10,20,50] are more precise than FA3[5,10,70]. The results of FA3[Poor] are clearly the worst. This flip angle combination not only leads to a 1.63% measurement bias, corresponding to about 22% overestimation of the global mean, but also a higher RPC (2.89%).

**Figure 2:**
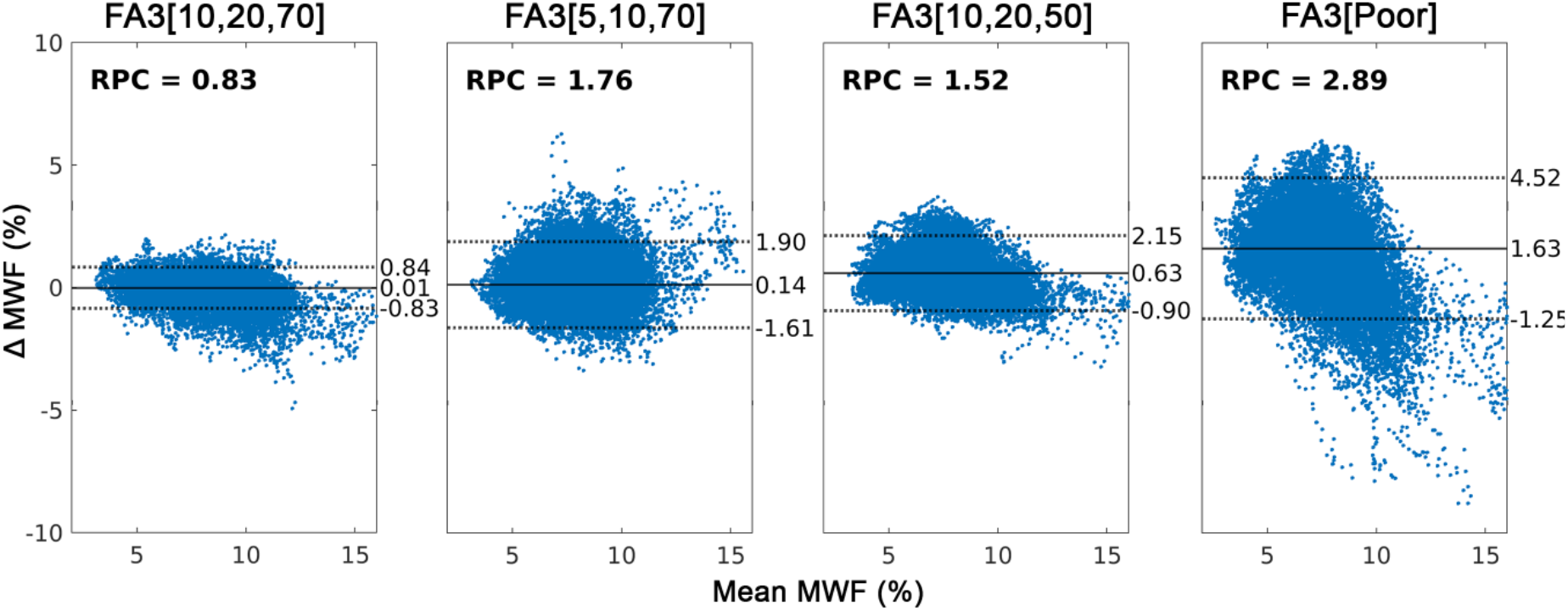
Bland-Altman analysis comparing MWF estimation from using data comprised 3 flip angles to 5 flip angles for 4 different flip angle combinations. Solid line represents the median and the dashed lines represent the 1.45IQR.

### 3.2. Test-retest analysis and comparison between protocols

The MWF maps of all protocols and all sessions for both MCR-MWI and MCR-DIMWI fittings from one of the subjects are shown in Figure 3. High-quality MWF maps can be observed from both models for all protocols. Highly myelinated fibre bundles such as the splenium of the corpus callosum and optic radiation have higher MWF values in respect of nearby tissues. Some inconsistent MWF estimation between the two sessions can, however, be observed in the frontal region even within the same protocol (see blue arrows).

**Figure 3:**
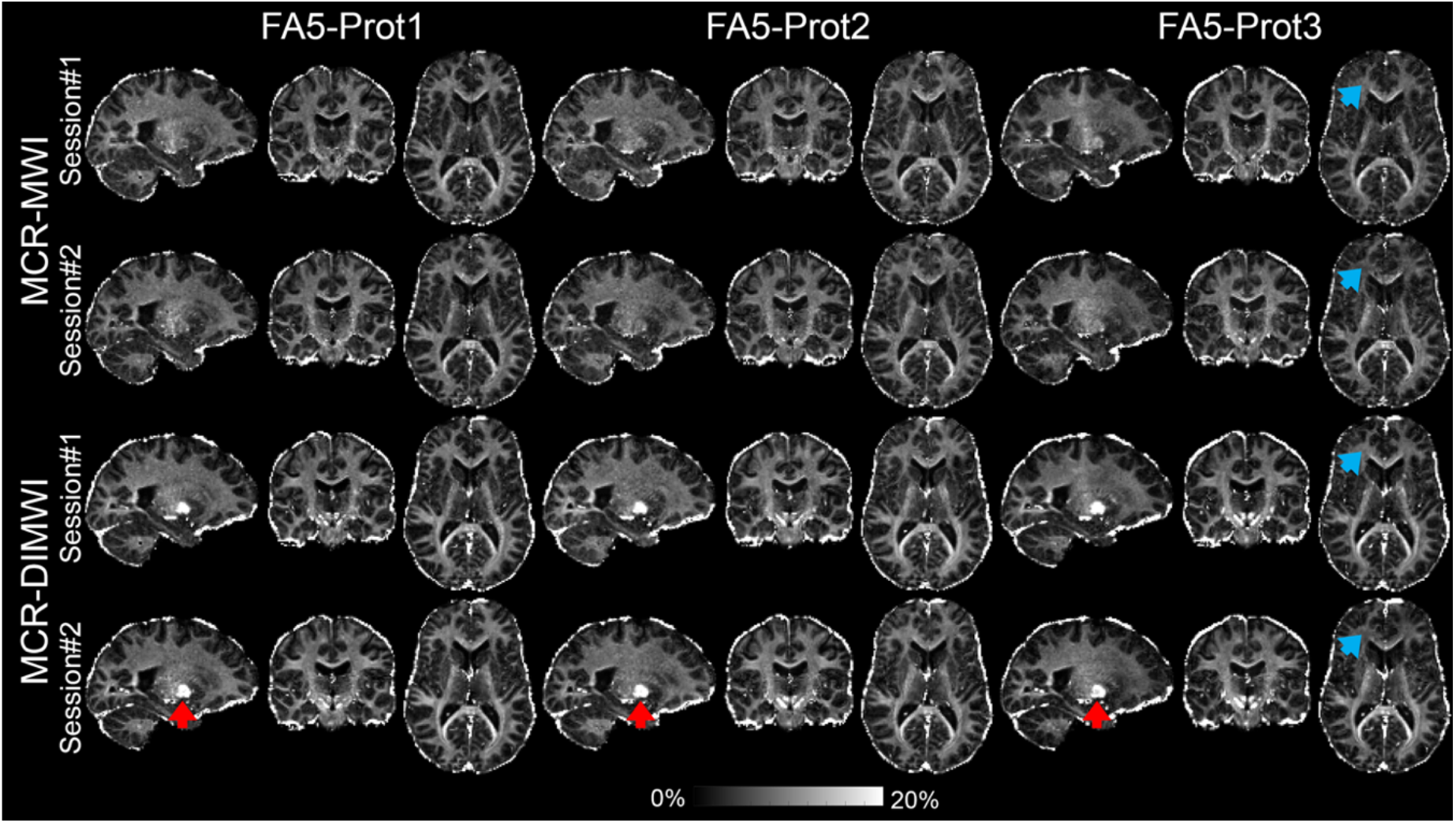
Example MWF maps on 1 subject. Both MCR-MWI and MCR-DIMWI produce high-quality MWF maps. Small contrast differences can be observed in the frontal regions, particularly between Protocol 3 and the other 2, which are potentially caused by the image artefacts in the data associated with different TRs (blue arrows). The MCR-DIMWI results have a slightly higher SNR appearance, despite the MWF values being incorrect in some deep GM nuclei (see red arrows) due to the fitting boundaries.

Figure 4 shows the cross-session Pearson’s R and CV of each protocol for each subject. Generally, the CVs are lower when the correlation coefficients are higher. No protocol-specific trend was observed at the subject level. One subject (Subject 3) shows a consistently lower correlation and higher CV than the other subjects which were attributed to the large respiration artefacts observed in the raw data (see Figure A.1).

**Figure 4:**
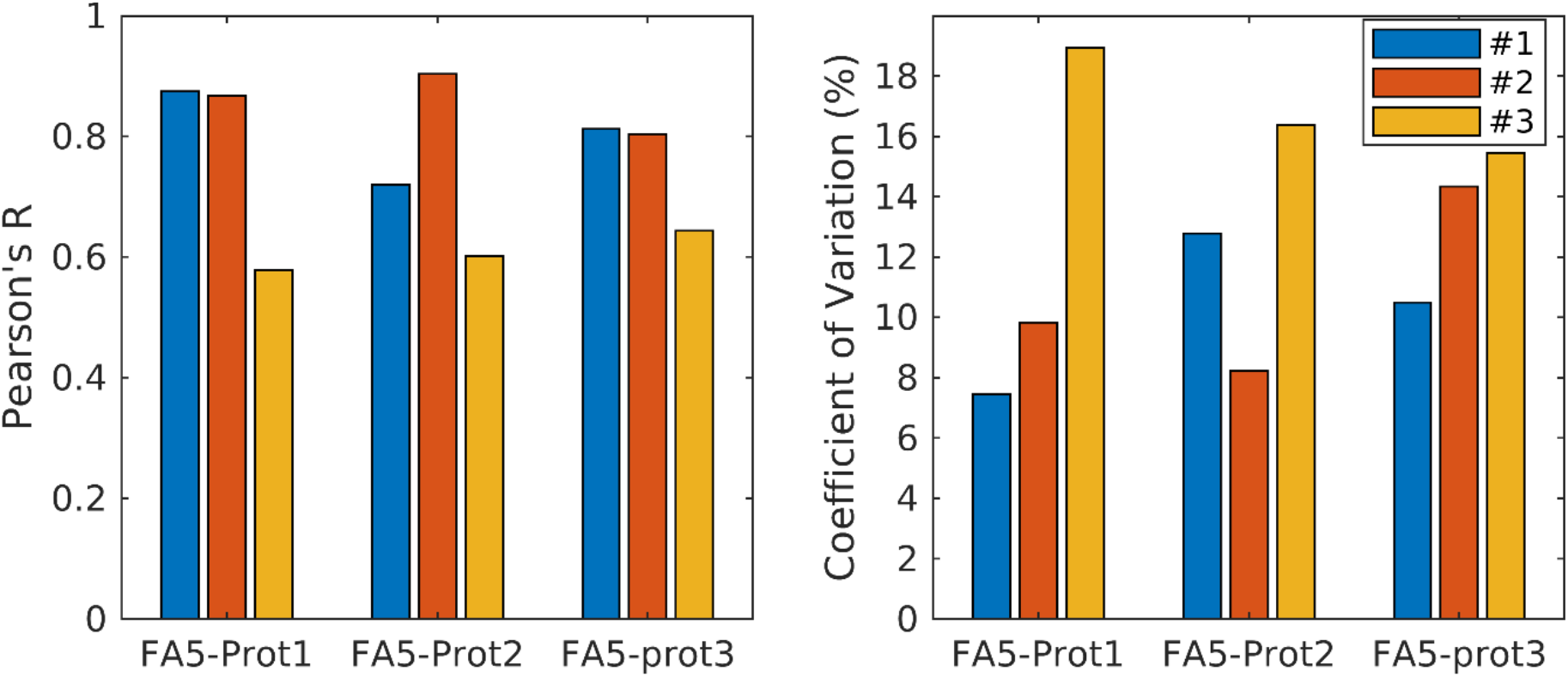
(Left) Cross-session Pearson’s correlation coefficients and (right) coefficients of variation at the subject levels (Colours represent subjects). Differences can be observed for the same GRE protocols across subjects. This illustrates the quality of the data has a substantial effect on the estimation reproducibility even with the updated weighting methods used (refer to Appendix A).

When pulling the results from all subjects together, the metrics should reflect more on the protocol-specific differences and be less affected by the quality of individual data. The differences in the protocol settings resulted in small MWF biases across the three protocols, as shown in Figure 5 (left panel), where a maximum bias of 0.70% MWF was observed between Protocols 1 and 3 (median MWF across all ROIs in GT = 8.5%) for using all 5 flip angles (0.96% for FA3[10,20,70]) with MCR-MWI. Test-retest analysis shows similar repeatability among the three protocols (R ≥ 0.7 in Figure 5, middle panel) and reducing the number of flip angles from 5 to 3 does not have a significant impact on the correlation coefficients between the two imaging sessions. A similar trend can also be observed in CV across the protocols (Figure 5, right) and reducing the amount of data leads to an increase in the CV which agrees with the simulations. The results of the same analysis on other tissue parameters can be found in Supplementary Figure S1. For other tissue parameters derived from Protocols 1, 2 and 3, the median ± IQR R_2,IW_* are 16.8 ± 3.1 s^-1^, 15.9 ± 2.2 s^-1^ and 16.4 ± 2.8 s^-1^; R_1,IEW_ are 0.63 ± 0.092 s^-1^, 0.67 ± 0.098 s^-1^, 0.73 ± 0.095 s^-1^; and k_IEWM_ are 3.3 ± 1.6 s^-1^, 2.6 ± 1.1 s^-1^ and 2.2 ± 1.0 s^-1^.

**Figure 5:**
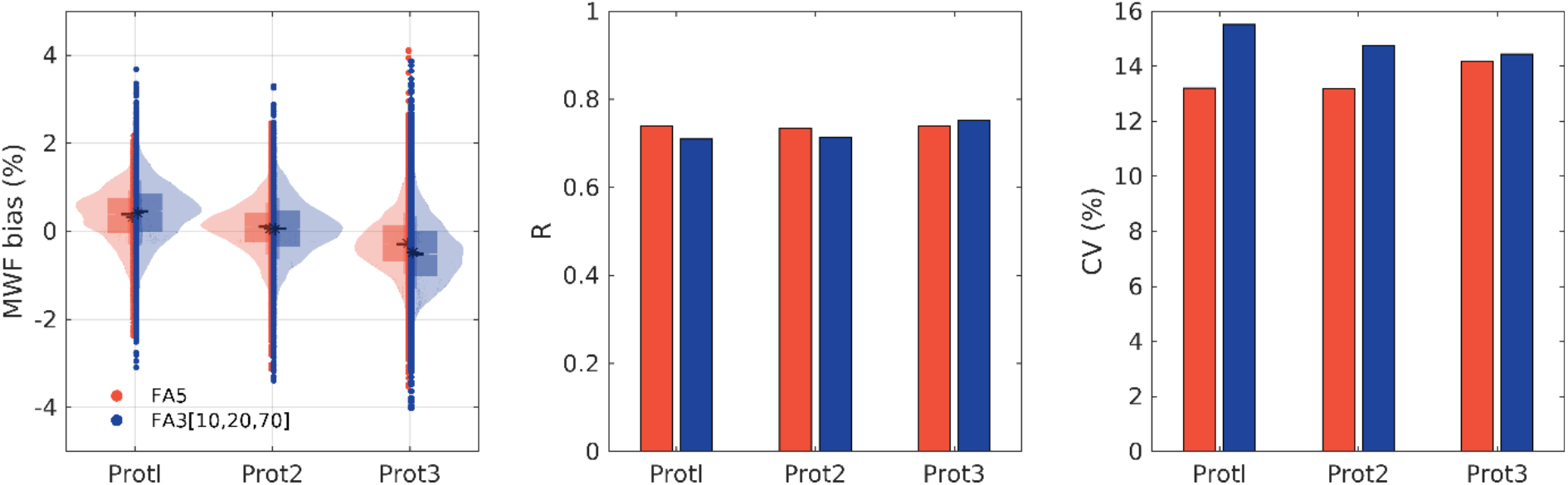
Protocol-specific differences based on MCR-MWI derived MWF at the group level. Red represents the results obtained from using all 5 flip angles while blue is derived from using 3 flip angles with the best combination. (Left) Boxplots of the MWF estimation bias. Box indicates the IQR; black line is the median and asterisk is the mean. (Middle) Pearson’s correlation coefficient and (right) coefficient of variation (CV) of MWF between the two sessions for all three protocols.

Between MCR-MWI and MCR-DIMWI, both show acceptable reliability (R ≥ 0.7 for MCR-MWI and R ≥ 0.8 for MCR-DIMWI; first and third columns in Figure 6) for all protocols and the metrics between protocols are comparable. For inter-protocol reproducibility, the correlation coefficients and CVs of MWF between different protocols are also in a similar range to the within-protocol results (second and fourth columns in Figure 6). In most cases, MCR-DIMWI improved both R and CV with respect to MCR-MWI.

**Figure 6:**
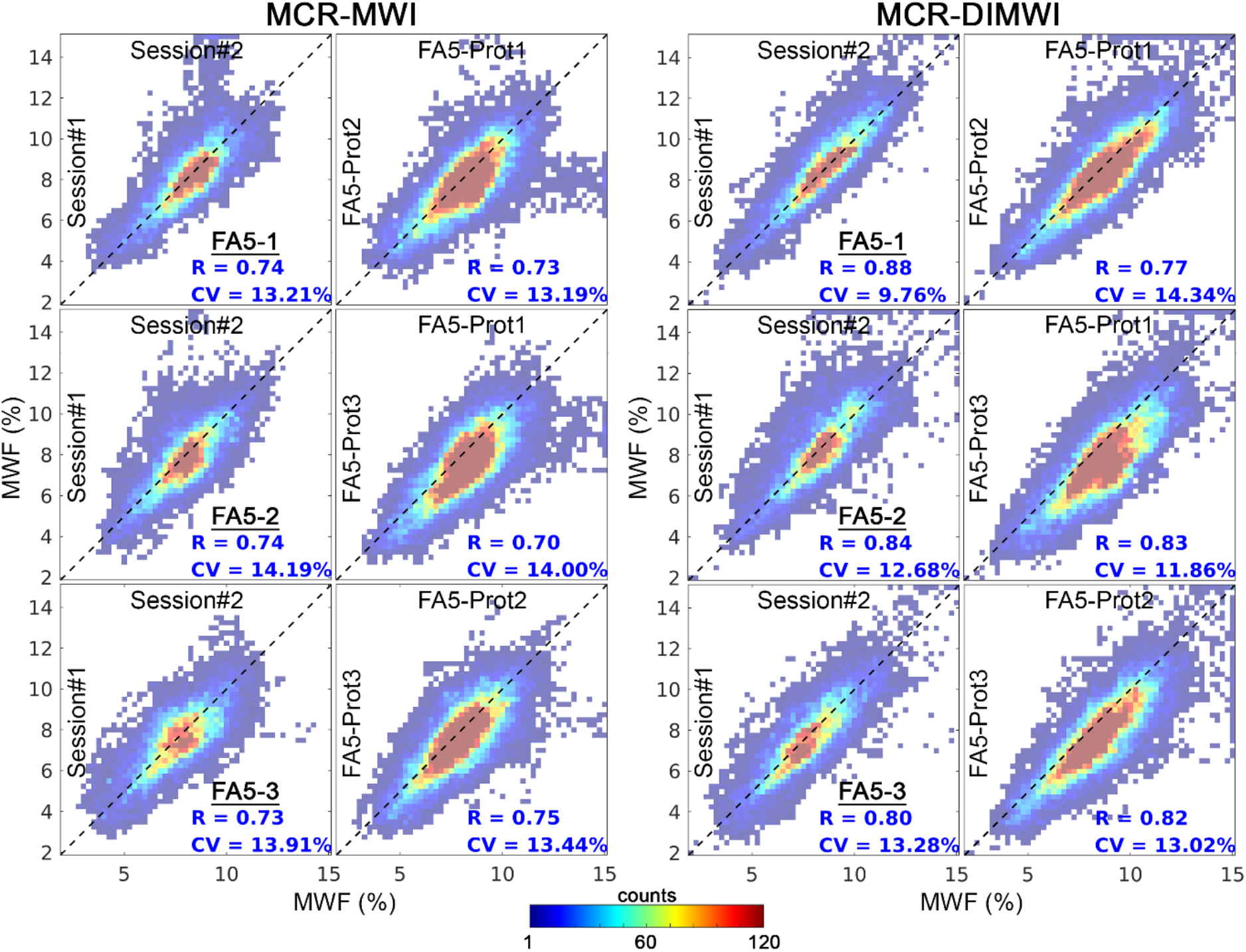
Test-retest repeatability analysis for the three protocols (first and third columns) and comparisons between protocols (second and fourth columns). Colours represent the number of the WM ROIs overlapped and the black dashed line is the unity line.

### 3.3. WM bundles-specific analysis

Eight major WM fibre bundles were selected to study the bundle-specific MWF profiles with Protocol 2 data (only one session). Among these bundles, only the ILF profiles from MCR-MWI are less similar among the 3 subjects. These differences between subjects could be attributed to the image artefact-induced error, bundle definition and coregistration accuracy in addition to individual differences. Similar shapes of the bundle profiles can also be observed between the left and right hemispheres of OR and ILF of each subject. The MCR-DIMWI profiles generally have greater similarity across subjects when compared to those from MCR-MWI, in terms of shape and values (see bottom panel of Figure 7), further demonstrating the greater robustness of this method in WM.

**Figure 7:**
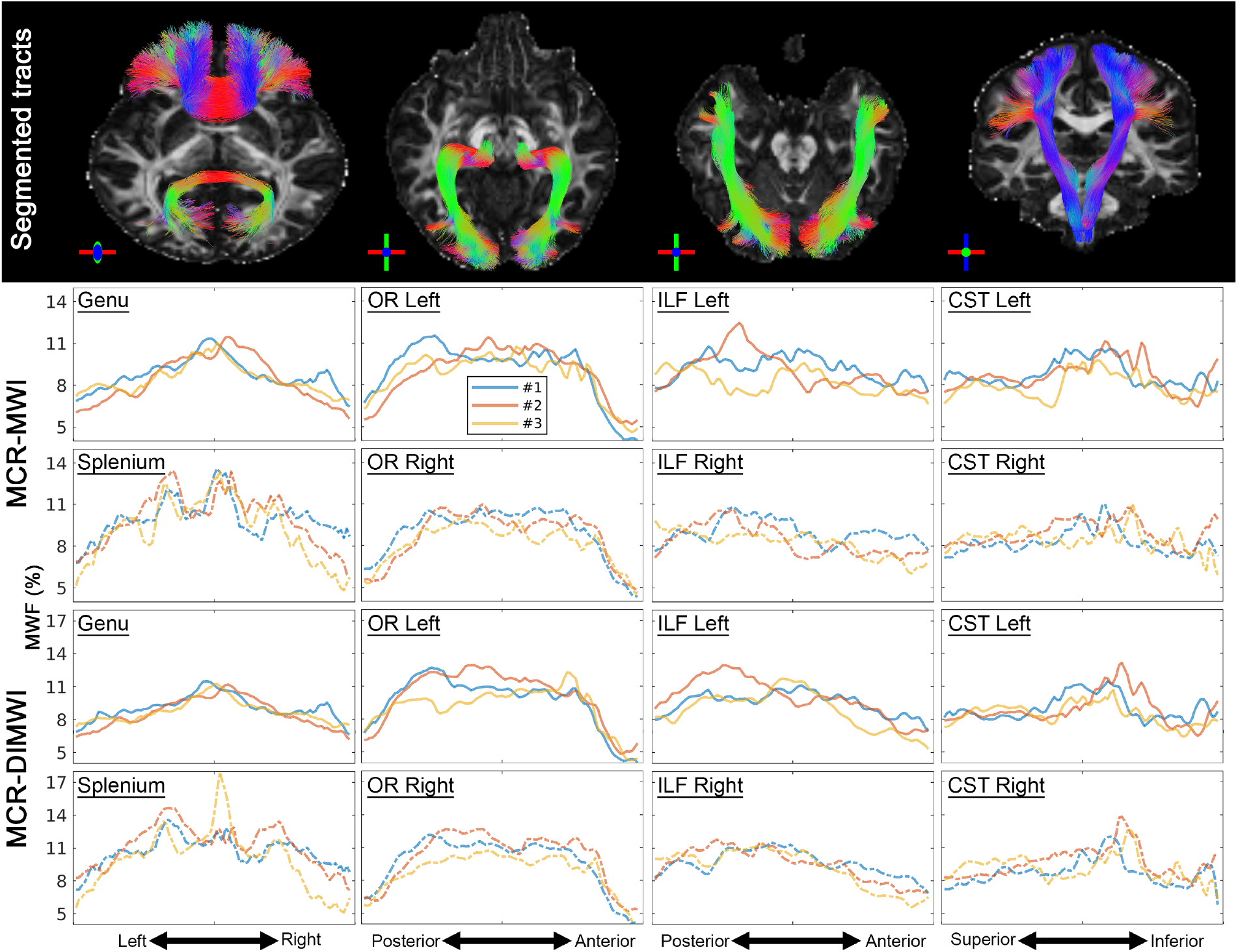
(Top row) Illustrations of representative WM bundle segmentation result on 1 subject overlaid over the DTI fractional anisotropy map. The middle panel (2^nd^ and 3^rd^ rows) shows MWF profiles obtained from Protocol 2 using the MCR-MWI model along the WM bundles shown on the top row (only one session is demonstrated). The 3 different subjects are represented by different colours. The bottom panel (4^th^ and 5^th^ row) shows MWF profiles obtained from the same protocol using the MCR-DIMWI model along the same WM bundles.

The bundle-specific profiles of 5 MCR-(DI)MWI derived tissue properties are shown in Figure 8 and the corresponding Pearson’s correlations between these parameters are shown in Figure 9. The MWF, k_IEWM_ and R_2,IW_* profiles derived from MCR-DIMWI are slightly, yet systematically, different from those obtained from MCR-MWI across all the studied bundles (see Figure 8 red vs blue lines). MCR-MWI seems to underestimate MWF and R_2,IW_*, while it overestimates g-ratio. These tissue properties show residual protocol dependence despite exhibiting similar profiles along with the WM bundles. Specifically, the short TR protocol (i.e., Protocol 1) has a higher MWF (and therefore a lower g-ratio), faster k_IEWM_ and slower R_1,IEW_ in contrast to the longer TR protocols (i.e., Protocols 2 and 3). Meanwhile, g-ratio profiles among all three protocols are comparable with values that are consistently higher than 0.8. Among these tissue parameters, weak correlations can be observed between MWF and k_IEWM_ (0.20) and R_2,IW_* (−0.33), and weak-to-moderate correlations among the relaxation and exchange parameters for MCR-MWI. These parameters become more linearly independent when diffusion-informed microstructure information is introduced and only moderate correlations are observed between R_1,IEW_ and R_2,IW_^*^ and between R_1,IEW_ and k_IEWM_ (as can be seen in Figure 9). Among all the examined parameters, MWF and g-ratio have a very high linear correlation for MCR-MWI (−0.93) but became moderate when DWI data is available (MCR-DIMWI: -0.53).

**Figure 8:**
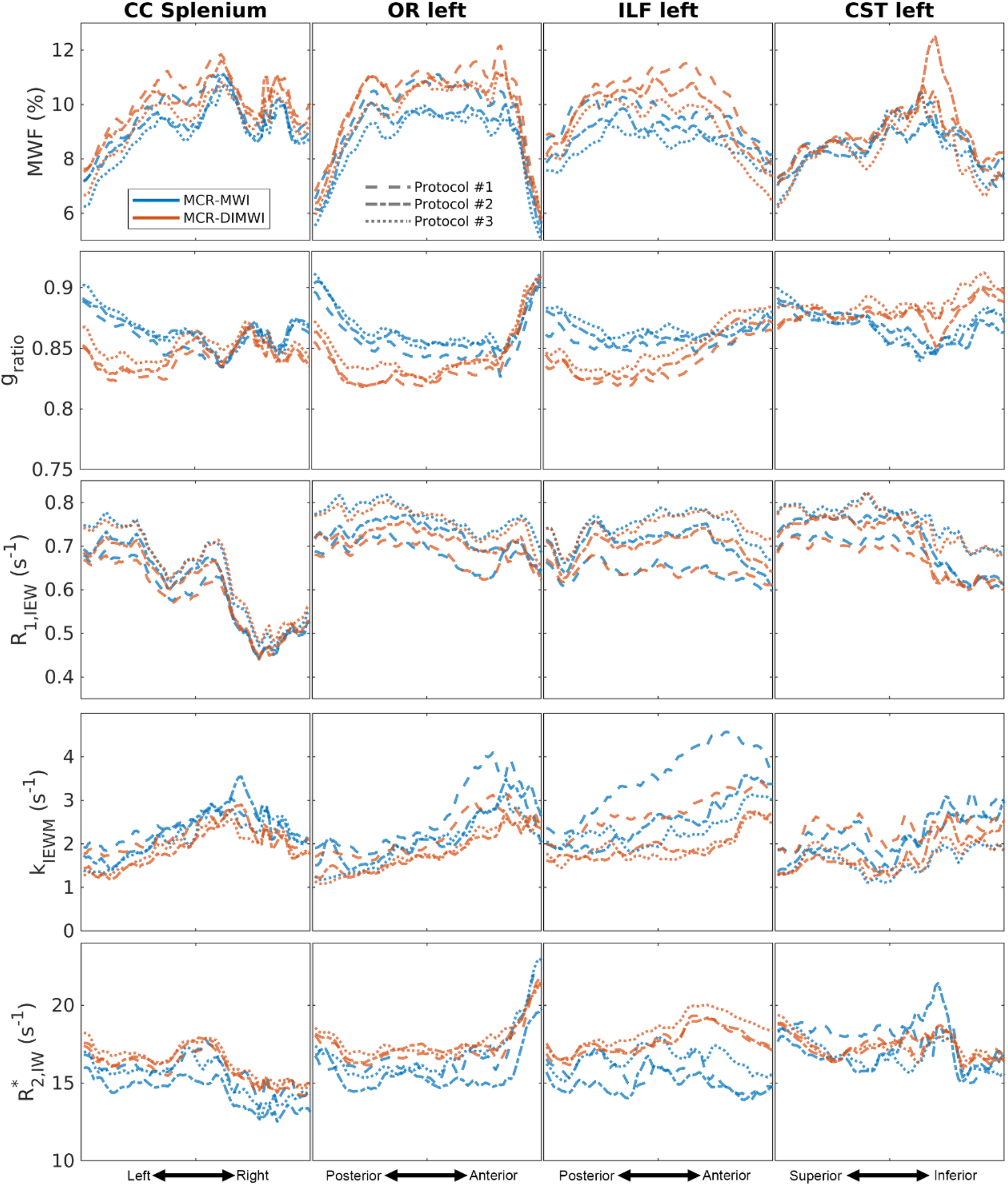
WM bundle profiles of MWF, g-ratio, R_1,IEW_, k_IEWM_ and R_2,IW_* averaged across the three subjects and the two imaging sessions. The models are represented by different colours while the line styles present the three GRE protocols.

**Figure 9:**
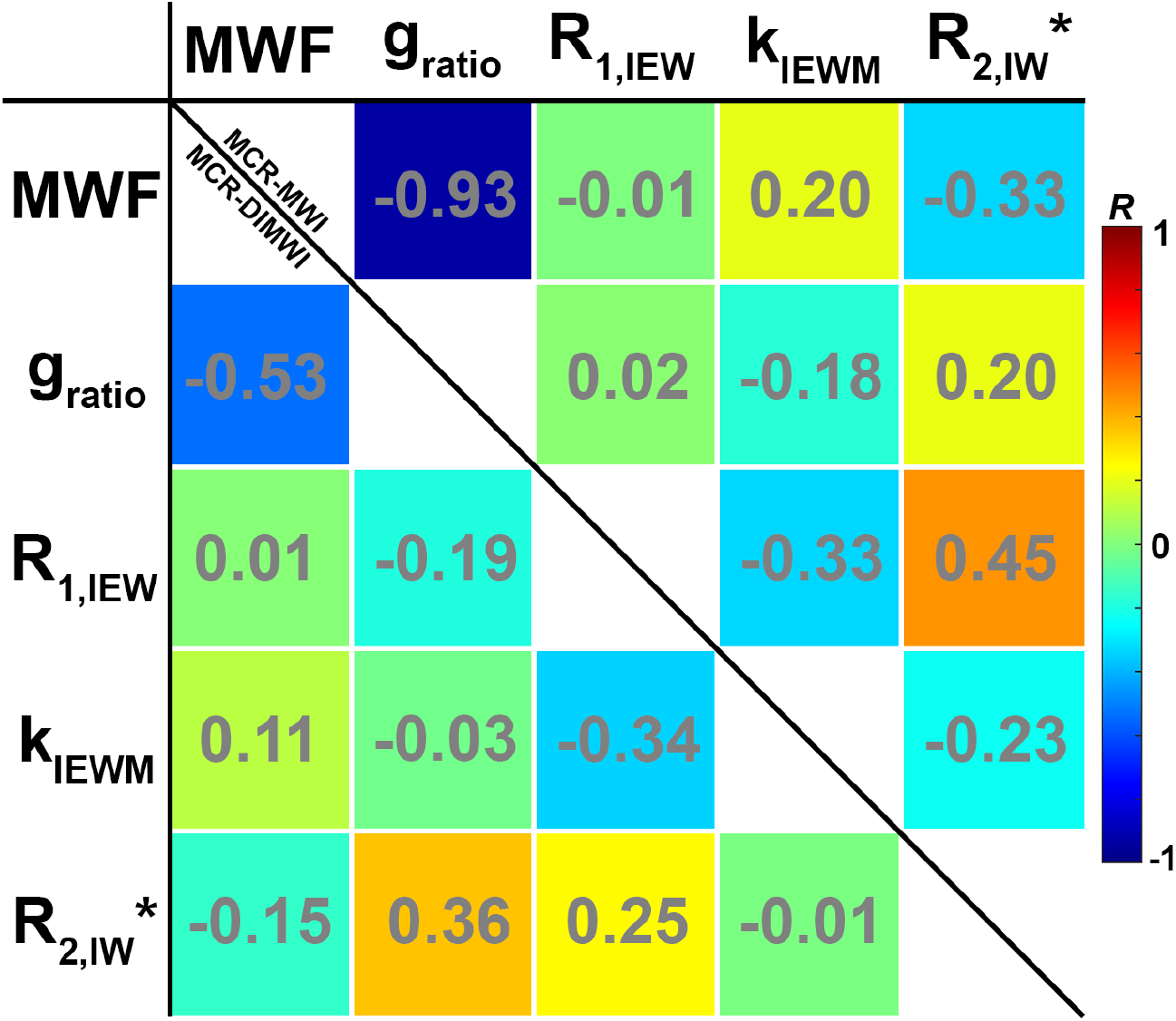
Pearson’s correlation coefficients between MWF, g-ratio, R_1,IEW_, k_IEWM_ and R_2,IW_* using the results of Protocol 2 on all ROIs. Upper triangle of the table are the results based on MCR-MWI while the lower triangle represents the results of MCR-DIMWI.

## 4. Discussion

In this work, we optimised the VFA acquisition scheme for MCR-MWI via numerical simulations. We observed that many 5 flip angle combinations were comparable to our initial protocol using 7 flip angles (Chan and Marques, 2020), albeit systematically having a minor penalty in precision because of the reduced measurement time (about 1.18-fold see Table 1). Furthermore, it suggested that MCR-MWI can be performed with as few as 3 flip angles if an appropriate flip angle combination is chosen. Hence, in this study, we acquired various protocols (different TR and TEs) based on a 5-flip-angle approach and confirmed that comparable MWF can be obtained using data from 5 flip angles and 3 flip angles via in vivo experiment. These findings support that MCR-MWI data acquisition can be performed in shorter and more clinically relevant scan time (between 8.5 min and 12 min at 1.5 mm isotropic resolution) without substantial degradation of the quantification result or utilisation of advanced image reconstruction techniques (Dong et al., 2021). Note that this outperforms some of the most recent advances on spin echo-based MWI using GRASE acquisitions with CAIPI acceleration (Piredda et al., 2021), where 1.6mm isotropic resolution was achieved in 10.5 min, and the MWF maps of MCR-MWI appear to be sharper than the results shown in (Piredda et al., 2021) (also see Figure S4).

We also evaluated the test-retest repeatability of MCR-(DI)MWI with 3 sets of GRE protocols. Good correlations (≥ 0.7) and acceptable CVs (< 15%) between the two imaging sessions are found in all protocols, suggesting MCR-(DI)MWI are reproducible methods for MWF quantification. These metrics will be useful for power analysis in future group/longitudinal studies. Despite the comparable correlations and CV observed in cross-protocol comparisons (see Figure 6), systematic MWF differences were found between the three GRE protocols (Figure 5). In particular, the protocol with the shortest TR (Protocol 1) generated higher MWF than the protocols with longer TR (Protocols 2 & 3). These systematic differences can also be found in both R_1,IEW_ and k_IEWM_. Previous studies have already shown that single compartment T_1_ mapping using the VFA method is susceptible to on resonance magnetisation transfer effect (Teixeira et al., 2019b, 2019a). For the shorter TR GRE sequence, the increased MT effect will result in stronger signal saturation and shorter apparent T_1_. In MCR-(DI)MWI, as the myelin water T_1_ is a fixed parameter, this effect will be compensated by having a higher myelin signal, fast R_1,IEW_ relaxation and/or faster exchange between the water compartments. While preliminary data suggests that balancing the MT effect across the flip angles of the GRE sequence with the same TR does not have a significant effect on the fitted parameters (data not shown), balancing the MT effect for different TRs could be important to be able to fully compare MWF in different implementations. Nevertheless, for the protocols explored here where the energy deposition varied by a maximum of 45% when going from Protocol 3 to Protocol 1, they only resulted in a change of 8.6% variation in MWF estimation. This change, when accounting for the large range of MWF values present in WM, is significantly smaller (4x) than what is observed for single compartment relaxometry performed on the same datasets (as shown in Supplementary material Figure S2.b).

The experiments of this test-retest study also revealed that the GRE image quality has a non-negligible impact on the quantitative results. As shown in Figure A.1, the artefacts are likely related to the physiological noise originating from respiratory cycles induced by B_0_ fluctuations (Moortele et al., 2002; Noll and Schneider, 1994), even in the absence of head movements, this can impact MWF computation. This is both supported by the similar degree of movement across flip angles between the subjects (derived from the image registration transformation matrices) and the fact that the artefacts become stronger at longer echo times. In most cases, it results in a signal reduction at longer echo times (the inconsistent phases spread the signal over the entire FOV), which is misinterpreted as having a higher concentration of myelin water (short T_2_* in MCR-(DI)MWI). This shortcoming is not specific to MCR-(DI)MWI but rather to all GRE analyses in general (Gelderen et al., 2007; Hu et al., 1995; Lee et al., 2006; Nam et al., 2015a; Wen et al., 2015). The presence of the artefacts causes regional MWF bias that results in inaccuracies even when the analysis is performed in large ROIs. This bias can be mitigated by using a weighted non-linear least square data fitting, utilising the signal amplitude to modulate the influence of the problematic data in later echoes (see Figure A.1). However, this introduces a small measurement bias across the brain, and the inconsistent repeatability metrics across different GRE protocols between the 3 subjects may still to some degree reflect the differences in data quality of the acquisitions (see Figure 4). A better solution would be using a navigator echo in the acquisition as suggested by (Nam et al., 2015a) and/or performing retrospective correction using field monitoring devices (Vannesjo et al., 2015) to minimise the physiology-induced B_0_ fluctuation and improve the reproducibility of MCR-(DI)MWI.

In addition to using the segmented fibre bundles for the test-retest analysis, we also took the opportunity to study the bundle-specific profiles of MWF on several WM structures. The advantage of ROI-based analysis is to detect a weaker effect, for example, progressive or early stage of demyelination onset, thanks to the averaging processes in anatomically and functionally equivalent areas as defined by automated WM segmentation tools based on the DWI data of an individual subject. This automated process is more reproducible than using manually defined ROIs, and it is more feasible to perform systematically across the whole brain. Good correspondence of MWF can generally be observed on the same bundles between the left and right hemispheres of the brain within the same subject, but there are also discrepancies, such as the magnitude (in CST) and shape (in ILF), between the subjects. Without any ground truth references and given the small sample size (only 3 in this study), interpretation of these subject-specific differences is less meaningful. Studying the WM bundle profile of microstructural parameters with a larger sample size would allow characterising the myelination of individual fibre bundles and establishing reference values to facilitate differentiation of healthy tissue from diseased tissue or myelination levels in populations with different cognitive performances. Alternatively, microstructural abnormalities based on MCR-(DI)MWI in individual subjects can also be performed using the “Detect framework” (Chamberland et al., 2021), and may provide complementary information to improve the detection accuracy of the DWI-derived microstructural parameters in which myelin information is not available.

In this context of performing fibre bundle-specific analysis, the additional cost of performing MCR-DIMWI is extremely low (note that the MCR-MWI’s VFA acquisition already has a shorter acquisition time than the DWI data despite being acquired at a higher spatial resolution), from which the microstructure parameters are more reproducible having the benefits of the additional information from DWI (Figure 6). Yet, MCR-DIMWI requires HCM together with the assumptions based on healthy WM tissue properties to reduce the number of fitting parameters. In disease tissue, the invalidation of these conditions can result in biased estimations. Further improvements in the MCR-DIMWI models such as incorporating realistic WM geometries (Hédouin et al., 2021) to relax the HCM assumptions or add more advanced compartmentalization frequency effects in the myelin water compartment (Hédouin et al., 2021; Yablonskiy and Sukstanskii, 2014) would be important to be able to better translate MCR-DIMWI to diseases and regions with high iron concentration (where we observed fitting error). These models that are likely to better characterize the real physical environment should maintain the reproducibility improvements provided by MCR-DIMWI while reducing the estimation bias that is currently observed between MCR-MWI and MCR-DIMWI.

MCR-(DI)MWI produces several other microstructural parameters, such as compartmental R_1_ and R_2_^*^, but they are not yet thoroughly studied. Pearson’s correlation analysis between these microstructural parameters on the 4 studied bundles indicates the low linear correlation within WM of these parameters to MWF, suggesting they may possess independent information signalling other differences in the microstructural environment other than myelin concentration, which should be the focus of future studies.

## 5. Conclusion

MCR-MWI is a reproducible measure for myelin quantification and incorporating DWI microstructural information can further improve the method’s reproducibility. Care should be taken when comparing MWF from GRE sequences with different repetition times and/or RF pulses of different power as the currently implemented models do not explicitly account for the unbalanced MT effects. We also demonstrate that it is possible to perform MCR-(DI)MWI with data from at least 3 flip angles without substantially degrading the measurement performance, allowing acquisition time to be reduced up to 40% of the original implementation and total acquisition times below 10 minutes with whole brain acquisition at 1.5mm isotropic resolution.

## Acknowledgement

This work was funded by the Netherlands Organisation for Scientific Research (NWO) with project number FOM-N-31/16PR1056. Maxime Chamberland was supported by the Radboud Excellence Initiative Fellowship.

## Appendix A

In the original implementation, the weighting term *w* in Eq. 4 was defined as:

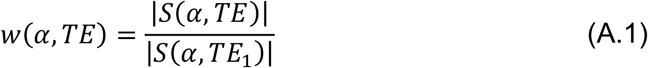

for each data point to account for the increasing physiological noise in later echoes (Chan and Marques, 2020; Nam et al., 2015b), corresponding to their own signal amplitude normalised by the magnitude of the first echo signal in the same acquisition. As illustrated in Figure A.1A, we observed significant data quality differences between the two sessions within the same protocol in the datasets of one of the subjects (see reduced R and increased coefficient of Subject 3 in Figure 4a and 4b). The image degradation increased with echo time, consistent with B_0_ field fluctuations related to respiratory motion (Moortele et al., 2002; Noll and Schneider, 1994), which manifests as an apparent signal dropout in the affected regions on later echoes and was mistaken as having high MWF in MCR-MWI (see Figure A.1B).

To minimize such artefacts in our test-retest analysis, we evaluated different signal weighting methods on the MCR-MWI fitting. Specifically, the weight *w* for a data point at TE and *α* was generalised to:

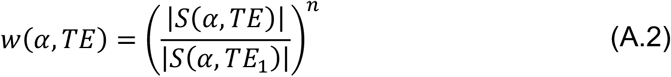

with n varying between 0 (standard RMSE), 1 (linear weighting found in MWF literature (Nam et al., 2015b)) and 2 (quadratic weighting) were tested on one subject with Protocol 1 data. As shown in Figure A.1B, when all data are equally weighted (i.e., n=0; left), there are visible differences on the MWF maps between the two sessions, for example, some WM regions show high MWF in Session 2 but not in Session 1 (see red arrows), which coincide with the observation that the VFA-mGRE data from Session 2 suffered more severe degradation. The MWF in these regions is gradually reduced with the increase of the exponential order n (Figure A.1B from left to right) and the contrast between the two sessions becomes more comparable.

The cross-session Bland-Altman plots of the MWF estimation using the TractSeg-derived WM ROIs from the three weighting methods show that the artefact-induced MWF bias results in a higher reproducible coefficient when the standard weighting approaches (i.e., n = 0 and n = 1) were used in the fitting, which can also be visually identified by the more scattering pattern in the plots. The quadratic weighting method significantly reduces the spreading, with the expense of a small estimation bias (0.44%) introduced in the process. If the data quality is relatively good for both sessions, the linear weighting method should be enough to minimize the differences introduced by image artefact and the quadratic weighting method would not provide further improvements as shown in Figure A.2.

**Figure A.1:**
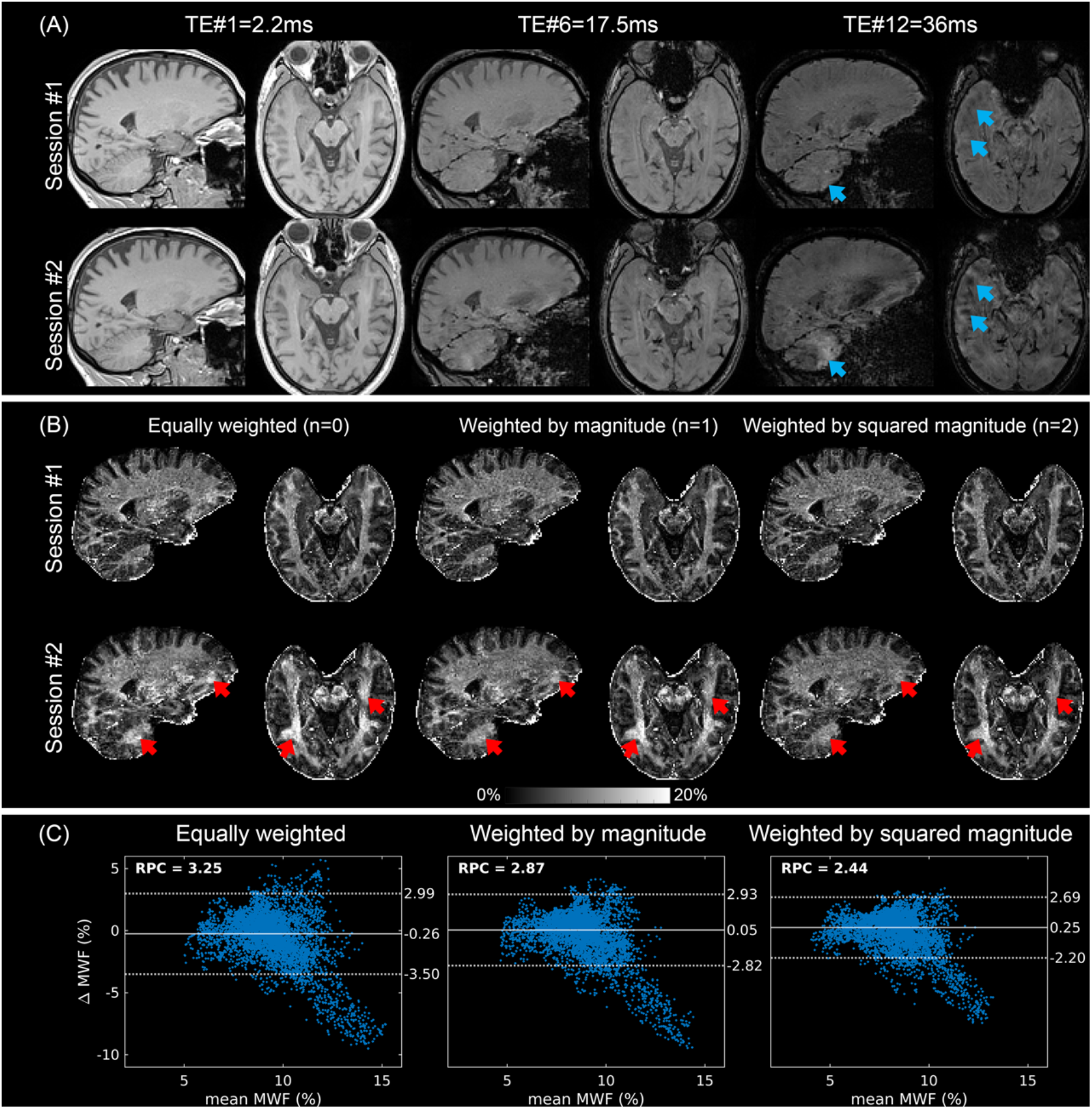
An illustration of the propagation of respiration-related artefacts to MWF maps and the impact of different data weighting strategies on 1 subject. (A) An example of the multi-echo GRE data from Protocol 1 with *α*=20° on 1 subject in two separate sessions. Blue arrows indicate some regions affected by the image artefact in the later echoes of the second session but not in the first session, though the data quality is very similar in the earlier echoes (rejecting the hypothesis of subject movement or cardiac noise). (B) The corresponding MWF maps when different weighting methods were used: (left, n = 0 in Eq. A.2) all data are equally weighted, (middle, n = 1) data are weighted by the magnitude of the signal, and (right, n = 2) data are weighted by the square of the magnitude signal. Red arrows show the estimation bias caused by the respiration artefact in the later echoes which are gradually reduced as the weighting power is increased. (C) Cross-session Bland-Altman plots of the MWF estimation in (B) using the TractSeg-derived WM ROIs. The artefact-induced MWF bias results in a higher reproducibility coefficient (RPC) when the fitting used standard weighting approaches (n = 0 and n = 1), which can be visually noticed by the increased distribution of the first two scatter plots. The quadratic weighting significantly decreases this, though a small estimation bias (0.44%) is introduced in the process.

**Figure A.2:**
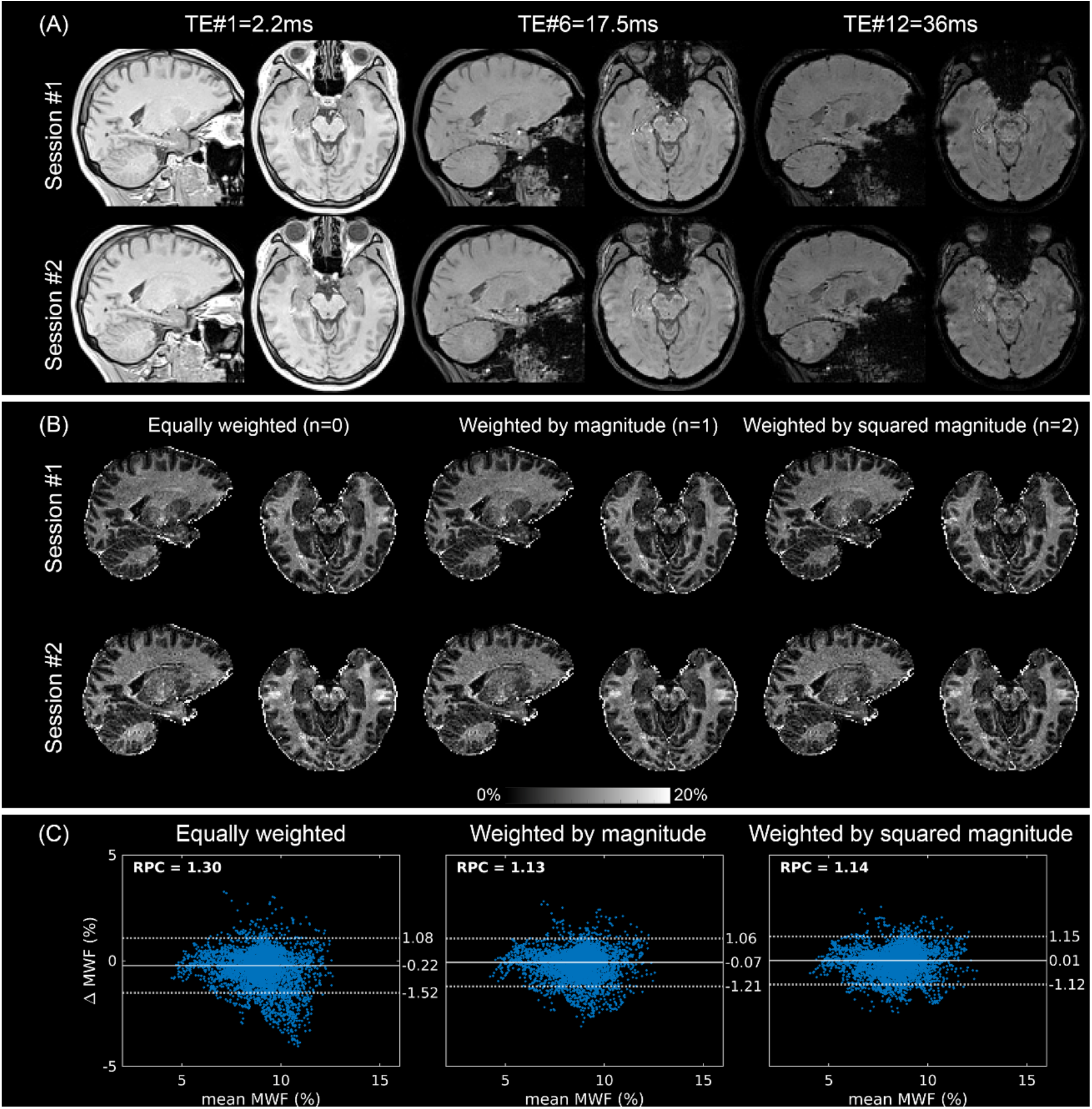
An illustration of the propagation of respiration-related artefacts to MWF maps and the impact of different data weighting strategies on another subject. In contrast to Figure A.1, the GRE data acquired in the two sessions show less significant artefacts. As a result, using linear weighting (n=1) or quadratic weighting (n=2) does not provide the improvements at the same management as shown in Figure A.1.

## Notes

[Funding] This work is part of the research programme with project number FOM-N-31/16PR1056 supported by the Netherlands Organisation for Scientific Research (NWO). Maxime Chamberland was supported by the Radboud Excellence Initiative Fellowship.

### Competing Interest Statement

The authors have declared no competing interest.

