## Supplementary for "On the performance of multi-compartment relaxometry for myelin water imaging – Intra-subject and inter-protocol reproducibility"

### Supplementary Material

#### 1. Tissue and acquisition parameters used in numerical simulations in Section 2.1

**Table S1:** White matter microstructure parameters used in numerical simulations. HCM: hollow cylinder fibre model.

| **Tissue properties** | **Value** |
|---|---|
| $S_0$ or $M_0$ (a.u.) | 10 |
| $g$ | 0.7[a] |
| $FVF$ | 0.73[a] |
| $\rho_{MW}$ | 0.42[b] |
| $\chi_I$ (ppm) | -0.1[c] |
| $\chi_A$ (ppm) | -0.1[c] |
| $E$ (ppm) | 0.02[c] |
| $T^*_{2,MW}$ (ms) | 10[d] |
| $T^*_{2,IW}$ (ms) | 64[d] |
| $T^*_{2,EW}$ (ms) | HCM |
| $\omega_{MW}$ | HCM |
| $\omega_{IW}$ | HCM |
| $\omega_{EW}$ | 0 |
| $T_{1,Myelin}$ (ms) | 234[e] |
| $T_{1,IEW}$ (ms) | 1050[f] |
| $k_{IEWM}$ ($s^{-1}$) | 2 |

[a](Stikov et al., 2015)
[b](Jung et al., 2018)
[c](Wharton and Bowtell, 2012)
[d](Nam et al., 2015)
[e](Du et al., 2014)
[f](Labadie et al., 2014)

**Table S2:** MGRE acquisition protocol used in numerical simulations.

| Parameters | VFA scheme |
|---|---|
| TE1(ms) | 2 |
| ΔTE (ms) | 3 |
| #TE | 12 |
| flip angle (°) | [5,10,20,30,40,50,70] |
| #Average | 1 |
| TR (ms) | 46 |
| $B_0$ (T) | 3 |

#### 2. Protocol-specific bias, Pearson's correlation and coefficient of variation on $R_{2,IW}$*, $R_{1,IEW}$ and $k_{IEWM}$

Figure S1: Protocol-specific differences based on MCR-MWI derived (a) $R_{2,IW}$*, (b) $R_{1,IEW}$, (c) $k_{IEWM}$ at the group level, using the same analysis as in Figure 5.

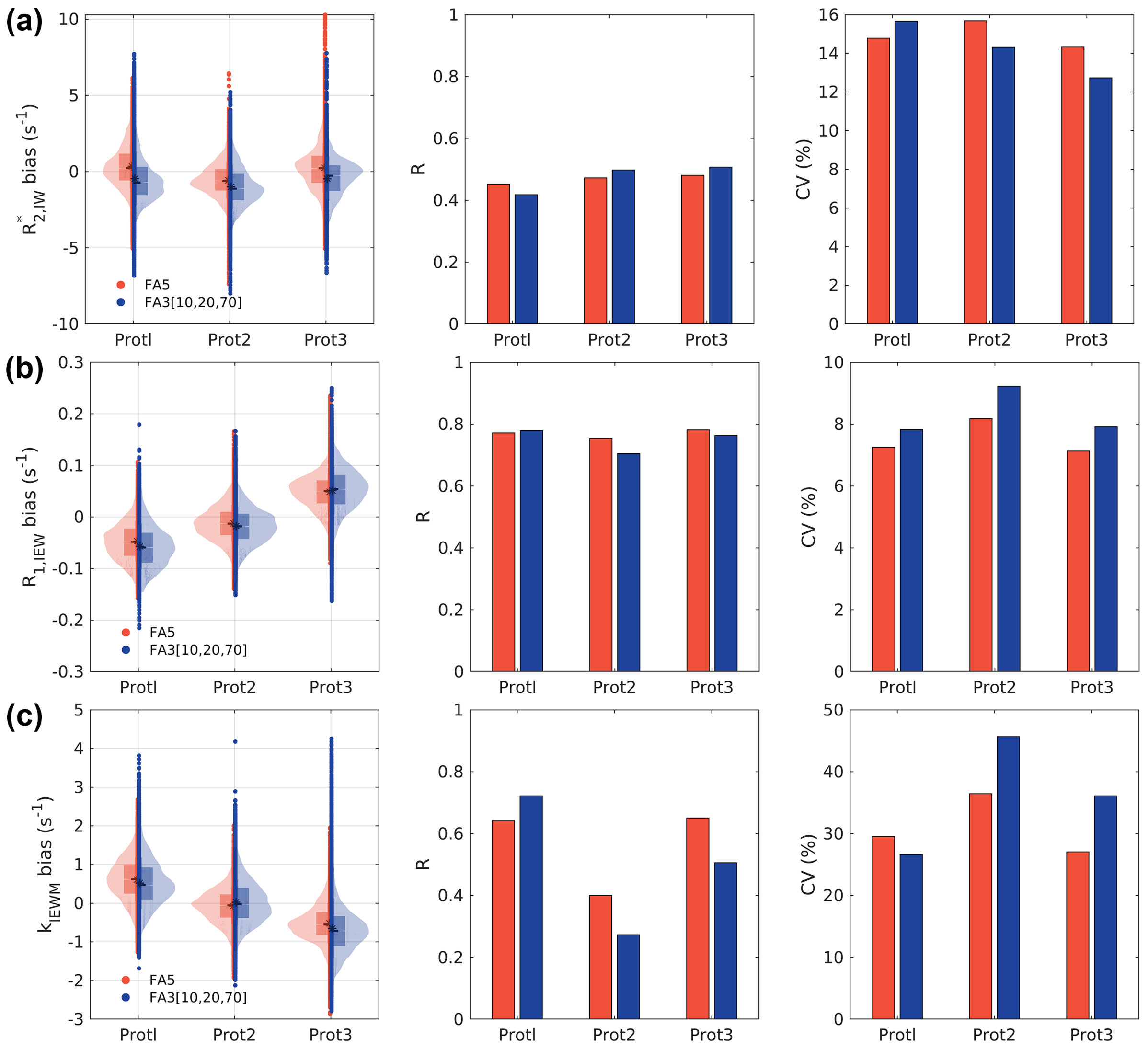

### 3. Protocol-specific single compartment $R_1$

Figure S2: (a) Protocol-specific single compartment $R_1$ bias using the same analysis as in Figure 5. (b) Normalised biases of MWF and single compartment $R_1$ by dividing the median by its own IQR of the derived biases across all ROIs. The relatively large bias of $R_1$ indicating that single compartment $R_1$ analysis is more sensitive to protocol-induced differences due to the narrow IQR in the estimation.

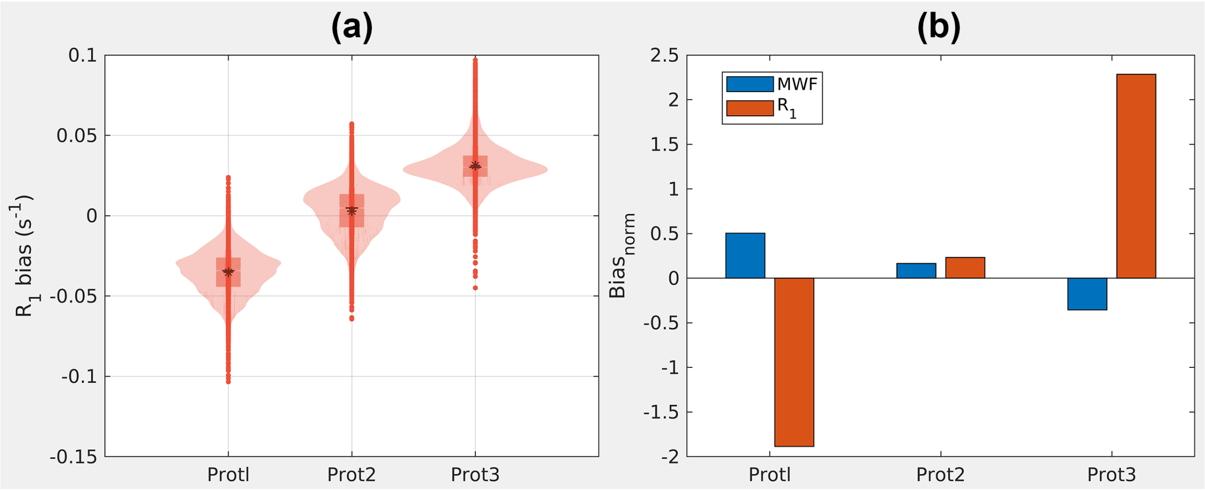

#### 4. Comparison when fitting the MCR-MWI signal model with and without EPG-X simulation

Figure S3: MWF differences induced by model consideration in various weighting methods mentioned in Appendix A. The data in red are derived using extended phase graph wth exchange (EPG-X) to simulate the steady-state signal while the results in blue are derived without using EPG (i.e., using the analytical solution of Bloch-McConnell equation). This result suggests that difference processing methods can also introduce systematic difference even using the same data.

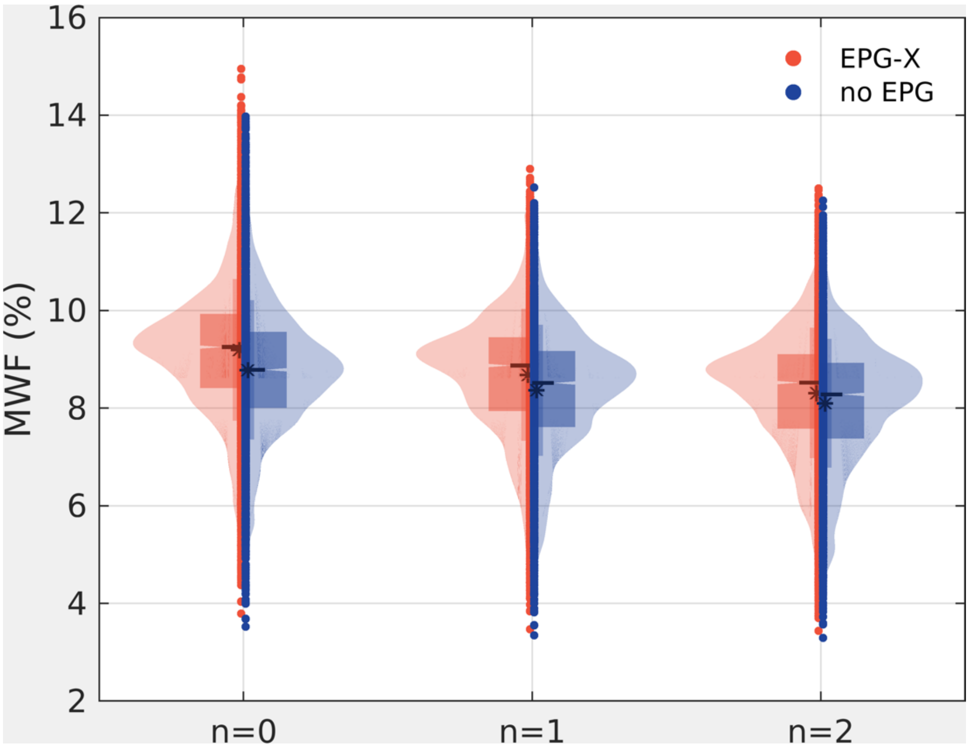

#### 5. MWF maps derived from data comprising 3 flip angles

Figure S4: MCR-MWI's MWF maps of all 3 subjects derived using only 3 flip angle of Protocol 1.

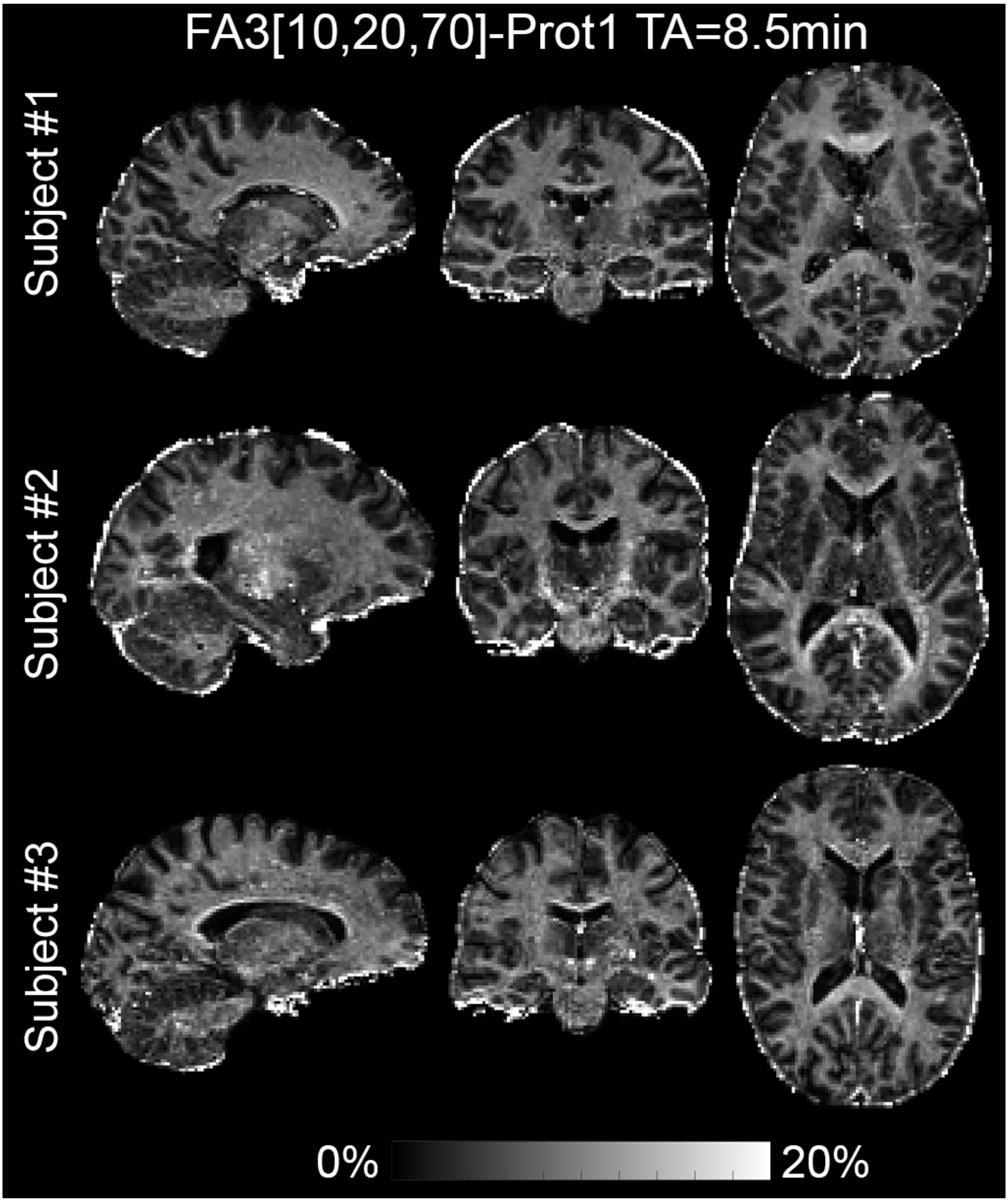

#### 6. Improving the splenium of corpus callosum segmentation from TractSeg

Figure S5: (a) Original TractSeg segmentation of the splenium of the corpus callosum and (b) the improved segmentation by manual annotation. When looking into the sub-components (c-e) of the original segmentation results on one subject, we observed that large irregularities of tract terminations within the fibre bundles and inconsistent with the results from other subjects, in turn, introducing non-structural related variations in the tract profiles. To minimise the effects of such discrepancies in the bundle-specific analysis, a filtering strategy was applied to target only the streamlines forming the conventional C-shape of the splenium of the corpus callosum by placing the filtering region of interests interactively at the sagittal stratum and occipital poles using FiberNavigator (Chamberland et al., 2014), resulting in only one component shown in (f). This process was repeated for all subjects to obtain comparable segmentation results.

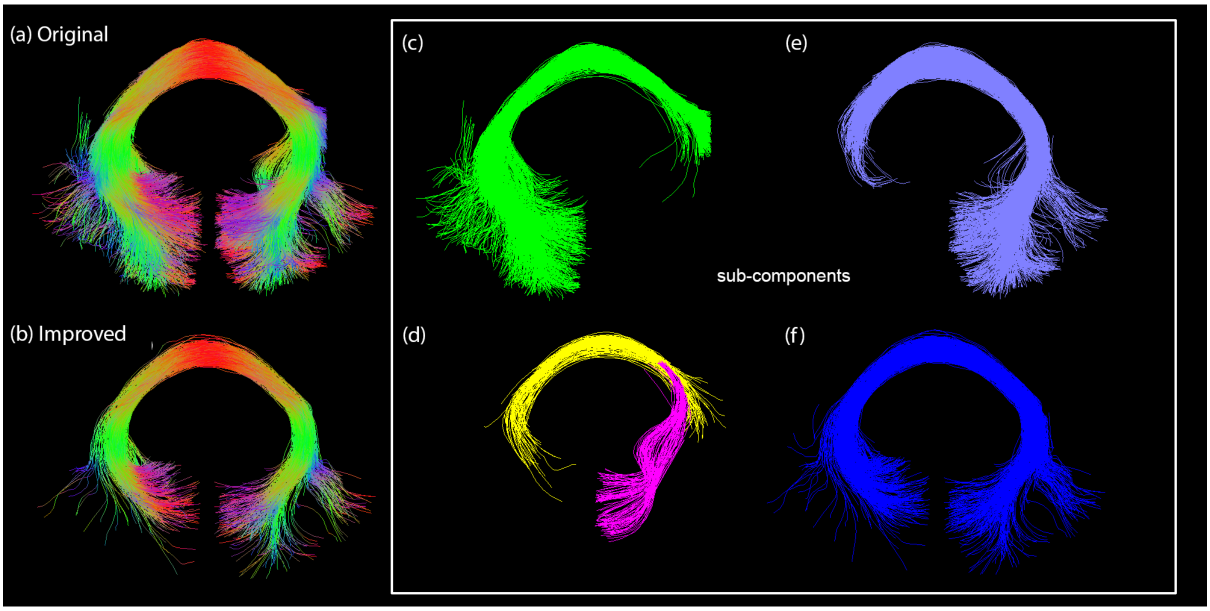
